# Metaproteomic Profiling of Fungal Gut Colonization in Gnotobiotic Mice

**DOI:** 10.1101/2020.12.24.424341

**Authors:** Veronika Kuchařová Pettersen, Antoine Dufour, Marie-Claire Arrieta

**Affiliations:** Department of Physiology & Pharmacology, University of Calgary, Calgary,Canada; Department of Pediatrics, University of Calgary, Calgary, Canada; International Microbiome Centre, Cumming School of Medicine, University of Calgary, Calgary, Canada; Department of Medical Biology, UiT The Arctic University of Norway, Tromsø, Norway

**Author notes:** Corresponding author Name: Marie-Claire Arrieta, Address: University of Calgary, Health Research Innovation Centre, 3330 Hospital Drive N.W., Calgary T2N 4N1, Alberta, Canada. Competing Interests statement: The authors declare no competing interest.

**Keywords:** Early Life Gut Microbiome, Quantitative Proteomics, Gut Fungi, Gnotobiotic Murine Model, Antimicrobials

## Abstract

Eukaryotic microbes can modulate mammalian host health and disease states, yet the molecular contribution of gut fungi remains nascent. We previously showed that mice exclusively colonized with fungi displayed increased sensitivity to allergic airway inflammation and fecal metabolite profiles similar to germ-free mice. To gain insights into the functional changes attributed to fungal colonization, we performed quantitative proteomic analyses of feces and small intestine of four-week-old gnotobiotic mice colonized with bacteria, fungi, or both. A comparison of fecal metaproteomic profiles between the mouse groups yielded broad changes in the relative levels of bacterial (46% of 2,860) and mouse (76% from 405) proteins. Many of the detected fungal proteins (3% of 1,492) have been previously reported as part of extracellular vesicles and having immunomodulating properties. Changes in the levels of mouse proteins derived from the jejunum (4% of 1,514) were mainly driven by proteins functional in lipid metabolism and apoptosis. Using metaproteomic profiling of gnotobiotic conditions, we show that fungal colonization profoundly impacts the host gut proteome. Our results suggest that an increased abundance of certain gut fungal species in early life may impact the developing intracellular balance of epithelial and immune cells.

## INTRODUCTION

The fungal portion of the mammalian gut microbiome, also referred to as the mycobiome, is estimated to constitute less than 0.1% of the gut ecosystem (1, 2). Although it has been significantly less characterized than the bacterial microbiome, the mycobiome has an important and often overlooked role in host health and disease (3-6). Cell numbers alone do not provide a comparable measure for the microbiome communities as fungal cells are up to 100 times bigger in volume and with genomes from 4 to 200 times larger than most bacteria (7), representing large biomass with potent production capacities for proteins and metabolites. Community ecology provides many examples of low abundance species with sizable impacts on community structure and function, showing that species’ importance cannot be predicted based upon their abundance in an ecosystem (8-10).

Fungi are an essential part of terrestrial and aquatic ecosystems, engaging in a wide variety of relationships with other members of microbial communities (*i*.*e*., bacteria, Archaea, viruses) and their plant and animal hosts (11-13). Bacteria modulate fungi’s ability to colonize mammalian hosts, as evidenced by clinical (14) and murine (15, 16) studies describing the effects of antibiotic treatment on the gut mycobiome. In general, antibiotic use promotes fungal overgrowth, and its immunological consequences can be detected at distant sites such as the lung (14, 16). Similarly, antifungals affect bacterial community structure and can have adverse effects on the host health (17). Controlled release of inbred laboratory mice into an outdoor enclosure further demonstrated the impact of fungi on the host immune functions (5). Although the mice rewilding enhanced the differentiation of immune cell populations previously associated with pathogen exposure, the resulting fungi-induced immune system changes occurred in the absence of an infection.

The inter- and intra-kingdom relations between microbes of the gut microbiome have been mostly described using the relative species abundance derived by DNA sequencing methods(18). However, this approach does not characterize molecular interactions between microbes that occur directly through secreted molecules and by physical contact, or indirectly by modulating the host immune response. Using a gnotobiotic mice model, we previously demonstrated the impact of gut fungal colonization on a defined bacterial consortium and the host (19). Gut fungi promoted shifts in bacterial microbiome ecology, and mice colonized exclusively with fungi displayed immunological features related to atopy in early life. Intriguingly, gut fungi had a marginal effect on the host fecal metabolome (19), suggesting that other functional mechanisms are at play. The results led us to investigate the host-microbiome interactions at the protein level, taking advantage of the defined experimental conditions and the availability of the bacterial species’ genomes.

Here, we show proteomic analyses of feces and small intestine of gnotobiotic mice, which were colonized with either 12 bacterial species (B), a group of 5 fungal/yeast species (Y), or a combination of the 17 fungi and bacteria (BY). We also evaluated the effect of microbiome perturbances to the metaproteome by analysing feces from mouse pups treated with antibiotic Augmentin (BY_ABX) or antifungal Fluconazole (BY_AFX). We observed dynamic relationships between the bacteria and fungi that were reflected by changes in the individual species proteome profiles derived from different colonization and treatment conditions. Metaproteomic analyses documented an extensive impact of microbial colonization on the host and revealed new features of the host protein response to fungal colonization. Proteomics of the jejunal tissue provided further insights on the host cellular pathways impacted by the microbial colonization and confirmed that gut fungi elicit distinct effects compared to bacteria.

## MATERIALS & METHODS

### Gnotobiotic Mice

Germ-free (GF) C57Bl/6J mice were obtained from and housed at the gnotobiotic mouse facility of the International Microbiome Centre (IMC) at the University of Calgary. Details of the animal experiments have been previously described (19), and all the animal work was conducted following animal protocols approved by the Institutional Animal Care and Use Committee. Briefly, female adult GF mice were orally gavaged twice, three days apart, with 100 μl of a consortium of microorganisms or kept under germ-free conditions. Colonization consisted of consortia of (i) 12 mouse-derived bacteria (20) (B), (ii) five yeast species previously linked to atopy and asthma risk in infants (21, 22) (Y), or (iii) a combination of all 17 bacterial and yeast species (B Y) (**Table S1**). Inocula were prepared under anaerobic conditions by mixing 100 μl of 2-day-old microbial cultures of each species. Bacteria were grown in fastidious anaerobe broth (LabM, Heywood, England, United Kingdom), and yeasts were grown in yeast-mold broth (YM; BD, Sparks, Maryland, USA). After the second gavage, mice were paired for mating on a 2:1 female:male ratio per cage. To ensure microbial colonization with the desired consortia in F1 mice, the corresponding inocula were further spread on the dams abdominal and nipple regions on days 3 and 5 after birth (23). Two groups of mice colonized with both bacteria and yeasts were treated with the antibiotic Augmentin (0.2 mg/ml; MilliporeSigma, St. Louis, Missouri, USA) or antifungal Fluconazole (0.5 mg/ml; MilliporeSigma) in sterile drinking water from day 7 to 14 after birth (groups BY_ABX and BY_AFX, respectively). Treatment solutions were prepared by dissolving the antimicrobials in distilled water, followed by filter sterilization. Mice were kept at maximum five animals per isolator cage and housed inside gnotobiotic flexible-film isolators, under a 12-h light/12-h dark cycle, 40% relative humidity, 22–25 °C, and ad libitum access to sterile food and water.

### Enrichment of Microbial Cells from Fecal Sample

Fecal samples were collected from gnotobiotic mice at the end of their 3^rd^ week of life and immediately stored at −80°C until use. After thawing at 4°C, pooled samples of ∼ 300 mg, originating from co-housed mice of the same treatment group, were subjected to differential centrifugation to enrich for microbial cells, according to previously described methodology (24). Briefly, each sample was resuspended in 4 ml of Phosphate-Buffered Saline (PBS), homogenized by using GentleMACS C tubes (Miltenyi Biotec, Bergisch Gladbach, Germany), and subjected to low-speed centrifugation at 20 × *g* for 5 min (Centrifuge 5810 R, Eppendorf, Hamburg, Germany) to eliminate gross particulate material. The supernatant was transferred to 50 ml conical centrifuge tube and kept at 4°C, whereas the pellet was resuspended in PBS. The washing step was repeated until the supernatant appeared translucent (5-7 times). The collected supernatant was centrifuged at 3,214 × g for 1 h, and the resulting pellet was subjected to cell lysis and protein digestion described below. A quality control step comprising of microscopic examination of Gram-stained fractions of the pellet was included to confirm bacterial and fungal cell extraction.

### Murine Intestinal Tissue Sample

Jejunal tissue dissected from the small intestine of 4-weeks old mice was cleaned of luminal debris with PBS and snap-frozen in liquid nitrogen and stored at −80°C until further processing.

### Protein Extraction

Enriched fecal microbiota and jejunal tissue samples were resuspended in lysis buffer [2% sodium dodecyl sulfate, 100 mM Triethylammonium bicarbonate (TEAB) buffer, 10 mM Ethylenediaminetetraacetic acid, and 1X Complete Mini EDTA free protease inhibitors (F. Hoffmann-La Roche AG, Basel, Switzerland), pH 8.0] in 1:4 w/v ratio and transfer into a 2 ml screw-cap tube with FastPrep Lysing matrix A (MP Biomedicals, Irvine, California, USA). Cells were mechanically disrupted by bead-beating for 2 × 3 min at 30 Hz (TissueLyser II, Qiagen, Hilden, Germany). The samples were incubated at −80°C for 10 min and at 95°C for 10 min, followed by centrifugation for 30 min at 18,000 x *g*, 4°C. To disrupt released cellular DNA that would interfere with downstream protein quantification, the supernatants were sonicated 3 x for 10s with 20s resting intervals on ice. Sonicated samples were centrifuged at 18,000 x *g*, 4°C, for 10 min, the supernatants collected, and protein concentration was measured by using DeNovix Spectrophotometer (DeNovix Inc., Wilmington, DE, USA).

### Sample Preparation for Label-free Quantitative (LFQ) Proteomics

The cell lysates of fecal microbiota-enriched samples were processed according to Filter-Aided Sample Preparation protocol (24, 25). Briefly, cell lysates containing 500 μg of total protein were incubated with 10 mM DTT in 100 mM Ammonium Bicarbonate (AmBic) at the solution to total protein ratio (v/w) 1:10 for 45 min at 56°C. The samples were then mixed with 8 M Urea in 10 mM HEPES pH 8.0 (UA) in YM-30 Microcon filter units (MilliporeSigma) and centrifuged at 10,000 × *g* for 15 min. After discarding the eluate, the filtration units were washed once with the UA buffer (10,000 × *g*, 15 min). Next, 50 mM iodoacetamide in UA was added to the filter, and samples were incubated in the dark for 20 min. The filter was washed twice with UA, followed by two washes with 50 mM AmBic. Proteins were digested with trypsin (Promega, Madison, Wisconsin, USA) in 40 mM AmBic at 37°C for 18 h (enzyme to protein ratio (v/w) of 1:100). The resulting peptide mixtures were desalted by using SepPak C18 solid-phase extraction cartridges (Waters, Mississauga, Ontario, Canada), lyophilized at 30°C in a vacuum concentrator (Speed Vac Plus SC110, Savant Instruments Inc., Holbrook, New York, USA) and stored at –80°C until further analysis. Prior to the LC-MS/MS analysis, the peptide mixtures were resuspended in 1% formic acid (FA). An aliquot of the tryptic digests was used to determine the concentration of the peptide mixtures by using Pierce™ Quantitative Colorimetric Peptide Assay (Thermo Fisher Scientific, Waltham, Massachusetts, USA).

### Tandem Mass Tag (TMT) Labelling

TMT samples were prepared according to the manufacturer’s instructions (TMTsixplex™ Isobaric Label Reagent Set, Thermo Fisher Scientific). Briefly, 150 μg of jejunal tissue cell lysates were reduced by incubating with 200 mM Tris (2-carboxyethyl) phosphine hydrochloride (TCEP) at 55°C for 1 h and alkylated with 375 mM iodoacetamide in the dark for 30 min. The proteins were precipitated by acetone, resuspended in 50 mM TEAB, and digested with trypsin at 37°C for 18 h. Resulting peptide mixtures originating from 4 treatment groups (GF, B, BY, Y) and 5 biological replicates (n=5) were labelled with TMT reagents distributed into 4 experimental groups (**Figure S1**). Each experiment had one TMT tag (131) that contained pooled samples, created by combining equal amounts of peptides (20 ug) of each sample from one experimental group. The pooled samples served as internal standards for normalizing the data across the experimental setups. For labelling, the peptides were incubated with TMT reagents for 1 h at room temperature. The reaction was quenched by adding 5% hydroxylamine, followed by incubation for 15 min. The resulting four multiplexed samples were desalted and quantified as described above.

### Mass Spectrometry Data Acquisition

Liquid Chromatography: Tryptic peptides were analysed on an Orbitrap Fusion Lumos Tribrid mass spectrometer (Thermo Fisher Scientific) operated with Xcalibur (version 4.0.21.10) and coupled to a Thermo Scientific Easy-nLC (nanoflow Liquid Chromatography) 1200 system. Tryptic peptides (2 μg) were loaded onto a C18 trap (75 um x 2 cm; Acclaim PepMap 100, P/N 164946; Thermo Fisher Scientific) at a flow rate of 2 ul/min of solvent A (0.1% FA and 3% acetonitrile in LC-MS grade water). Peptides were eluted using a 120 min gradient from 5 to 40% (5% to 28% in 105 min followed by an increase to 40% B in 15 min) of solvent B (0.1% FA in 80% LC-MS grade acetonitrile) at a flow rate of 0.3 μL/min and separated on a C18 analytical column (75 um x 50 cm; PepMap RSLC C18; P/N ES803; Thermo Fisher Scientific).

LFQ: Peptides were electrosprayed using 2.1 kV voltage into the ion transfer tube (300°C) of the Orbitrap Lumos operating in positive mode. The Orbitrap first performed a full MS scan at a resolution of 120,000 FWHM to detect the precursor ion having m/z between 375 and 1,575 and a +2 to +7 charge. The Orbitrap automatic gain control (AGC) and the maximum injection time were set at 4 x 10^5^ and 50 ms, respectively. The Orbitrap was operated using the top speed mode with a 3 sec cycle time for precursor selection. The most intense precursor ions presenting a peptidic isotopic profile and having an intensity threshold of at least 5,000 were isolated using the quadrupole and fragmented by higher-energy collisional dissociation (HCD, 30% collision energy) in the ion routing multipole. The fragment ions (MS2) were analysed in the ion trap at a rapid scan rate. AGC and the maximum injection time were set at 1 x 10^4^ and 35 ms, respectively, for the ion trap. Dynamic exclusion was enabled for 45 sec to avoid the acquisition of the same precursor ion having a similar m/z (plus or minus 10 ppm).

MS3-TMT: Analysis of TMT labelled peptide mixtures was carried out on the Orbitrap Fusion Lumos Tribrid mass spectrometer (Thermo Scientific) (control software Xcalibur™, version 4.0.21.10) using a data-dependent method with multi-notch synchronous precursor selection MS3 scanning for TMT tags as described previously1. The Orbitrap was operated with a positive ion spray voltage of 2.1kV and a transfer tube temperature of 300C. The scan sequence began with an MS1 spectrum (Orbitrap analysis, resolution 120,000; 375–1,575 m/z, AGC target 1e4, maximum injection time 50 ms). The maximum number of precursors within a 3 sec cycle time were fragmented using of collision-induced dissociation (normalized collision energy 35%) and the fragmented ions were analysed in the ion trap (turbo mode; maximum injection time 50 ms). To quantify the TMT reporter ion, we collected MS3 spectra using a method in which the top 10 MS2 ions were fragmented using HCD (collision energy 65%). The MS3 were analysed in the Orbitrap (AGC 1 x 10^5^; maximum injection time 105 ms; isolation window 2 m/z; resolution 50,000; scan range m/z 100-500). All acquisition methods are available in a summary format, as Supplementary F**igures S2** (LFQ) and **S3** (TMT MS3).

### Protein Identification and Quantitation

The raw spectral data were processed using MaxQuant (26) (version 1.6.5.0). For the LFQ data, the Andromeda search engine (27) integrated into the MaxQuant framework performed the spectral data search against a matched protein database (**Table S1**) composed of 12 genome-derived proteomes for the bacterial species (downloaded 26^th^ November 2018 from NCBI) and 6 UniProtKB-derived protein databases for 5 fungal species and mouse (downloaded 30^th^ December 2018). The MS3 spectra were searched against the mouse database.

For the MS data search, enzyme specificity was set to trypsin, allowing N-terminal cleavage to proline, and for ≤2 missed tryptic cleavages. Default settings were used for the MaxQuant searches, except that lysine acetylation and glutamate/glutamine conversion to pyroglutamic acid were set as variable modifications. N-terminal acetylation, methionine oxidation, and carbamidomethylation of cysteines was set as fixed modifications. The initial allowed mass deviation of the precursor ion was set to ≤20 ppm, and the allowed value for the fragment mass was set to ≤0.5 Da. Match between runs was used, with match time window 0.7 min and alignment window time 10 min. The maximum false discovery rates (FDR) at peptide and protein levels were set to 1%. Proteins LFQ intensities were determined by using the MaxLFQ algorithms(28). TMT-MS3 data were processed using MaxQuant using Reporter ion MS3 and TMT6plex-Lys126-131 internal labels, with the search setting described above.

### Proteins Functional Analyses

The MaxQuant output data were analysed by using Perseus (version 1.5.6.0) (38). Annotations for the identified proteins were downloaded from the UniProtKB database (29). Protein functional analyses, including metabolic pathway analysis, were performed using the DAVID (30) and STRING-db (31) tools.

### Experimental Design and Statistical Rationale

We performed LC-MS/MS analyses from 3-4 replicates of pooled fecal samples collected from mice belonging to each treatment group that underwent the same microbial colonization and was housed in the same cage and gnotobiotic isolator. Jejunal tissue samples were derived from 20 animals (5 GF, 5 B, 5 BY, and 5 Y mice), and the TMT experimental design is described in **Figure S1**. All samples were analysed by MS in a random order, and only proteins identified in at least two biological replicates were included in the analyses. To assess the biological variability of each experimental group, we calculated the Pearson correlation coefficients based on the protein intensities of each sample (**Table S2**). Before a normalization step and statistical analyses, the proteins LFQ intensities were log_2_-transformed. To correct for differences in the sample amounts injected into LC-MS/MS, we normalized the relative protein amounts by dividing each protein LFQ intensity by the median intensity for all proteins in a given replicate (32). To account for potential peptide loading differences in fractions of the 4 different 6-plex TMT batches, we applied a correction factor based on the pooled samples containing TMT^6^ label 131 (**Figure S1**). The TMT spectra intensities were first median normalized as described above and then each relative protein amount was divided by the correction factor.

To identify proteins with levels that differ substantially among the strains, we performed analysis of variance (ANOVA) to compare the global mean level of each protein against its corresponding amount in each condition. For the ANOVA test, FDR calculations were performed in Perseus by a permutation-based procedure with 250 randomizations and a cut-off of 5%. To determine the exact pairwise differences in protein levels, Tukey’s honestly significant difference (THSD) was performed on ANOVA-defined significant hits. Principal component analysis (PCA) was performed by using MetaboanalystR 3.0 (33), using log-transformed data normalized data by median. Hierarchical clustering analysis based on the protein levels was performed in Perseus, by using Euclidean distance, average linkage, no constraints, and pre-processing with k-means. Data are available via ProteomeXchange with identifier PXD019355.

Data Accession: PXD019355

Name: Metaproteomic Profiling of Functional Changes Driven by Gut Fungal Colonization

Username

Password: 2DSZcyWZ

## RESULTS

### Fecal metaproteome of defined gnotobiotic model

Using shotgun proteomics, we profiled the fecal metaproteomes of germ-free mice (GF), and mice colonized with either 12 bacterial species (B), five fungal species (Y), or both bacteria and fungi (BY) (**Table S1**). We enriched microbial cells from feces of four-week-old mice by differential centrifugation, digested cell lysates by trypsin, and analysed the resulting peptide mixtures using LC-MS/MS (**Figure 1A**). We queried the acquired mass spectra (5.3 x 10^6^ MS/MS) against a combined protein database of the 17 microbial species and mouse (**Table S1**). The proteomic search identified 70,190 unique peptide sequences (**Table S3**), mapped to 6,675 proteins (**Table S4**). Of these, 68% were bacterial, 22 % fungal, and 10% were mouse proteins (**Figure 1B**). Up to 58% of all detected proteins were quantified using label-free quantification. Principal component analysis based on the protein relative levels identified three main clusters (**Figure 1C**), suggesting similar protein profiles of Y and GF, B and BY, and BY mice groups treated with antimicrobials (BY_ABX and BY_AFX).

**Figure 1.**
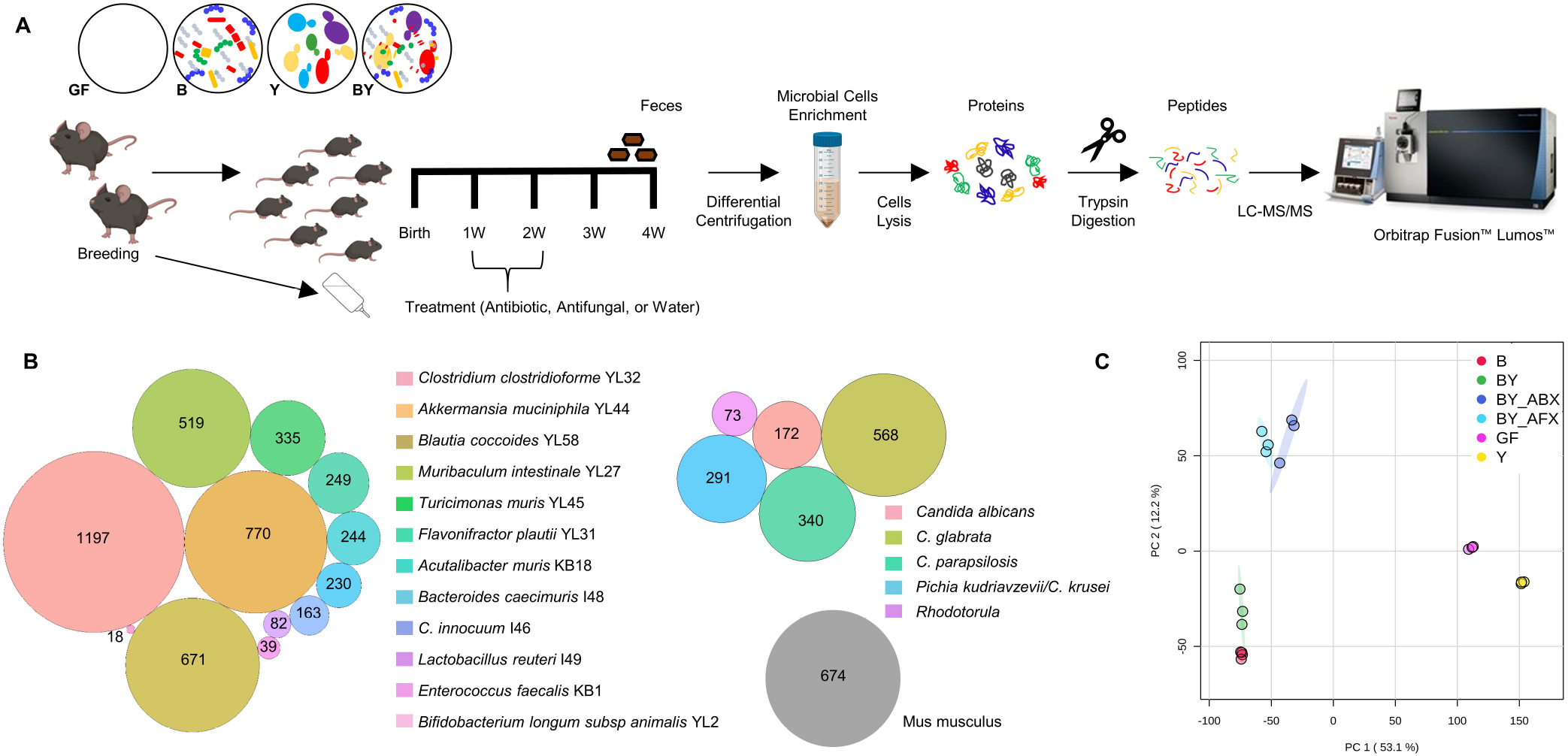
Description of a gnotobiotic colonization model by label-free quantitative metaproteomics. A) Experimental design of shotgun label-free proteomic experiment, including enrichment of fecal microbial cells by differential centrifugation. B) Detected fecal metaproteome composed of proteins originating from 12 bacterial, 5 fungal species, and mouse. C) PCA score plot based on the relative levels of 3,845 proteins quantified for mouse, bacteria, and fungi/yeast. X- and Y-axis show the first and second principal components, accounting for 53.1% and 12.2% of the total variation, respectively. Abbreviations of the mice treatment groups: B, bacteria; BY, bacteria-yeast; BY_ABX/AFX, bacteria-yeast and antibiotic or antifungal treatment; GF, germ-free; Y, fungi/yeast.

From the bacterial proteomes, 4.4% of detected proteins shared peptides with other protein groups (**Table S5**). However, the vast majority of the identified peptides belonged to a single protein group (leading protein group); that is, the sequence coverage of peptides unique for the leading protein group was considerably higher than the sequence coverage of peptides belonging also to alternative protein groups. For the fungal proteomes, a high percentage of identified peptides belonged to multiple homologous proteins from different strains of the same species (*e*.*g*., *Candida albicans* strains SC5314, WO-1, CD36) (**Tables S5**). This was due to the use of UniProtKB reference protein databases for the fungal species, which did not have sequenced genomes, contrary to the bacterial strains (34). The shortcomings of using not perfectly matched protein databases were most apparent for two fungal strains (*Pichia kudriavzevii* and *C. krusei*), which were recently re-classified as strains of the same species (35) and did not have available individual databases. For these fungi, we used a common database based on reference *P. kudriavzevii* strains available in UniProtKB, resulting in decreased specificity of the MS data searches.

Using genome-based protein databases for the bacterial stains also had the advantage of accurate assignment of proteins for the 12 different bacterial strains, for which we did not observe any species cross identifications. Nevertheless, the general UniProtKB databases did not compromise the accuracy of fungal and mouse protein identification, and we observed minimal cross-kingdom identifications (*e*.*g*., same peptides mapped to a fungal and mouse protein), with these proteins being manually filtered out.

### Inter-kingdom interactions influence the proteome response of gut bacterial species to antimicrobials

From 12 bacterial strains used for the gnotobiotic mice colonization, *Clostridium clostridioforme* YL32 had the most detected proteins (**Figure 1B**), while *Akkermansia muciniphila* YL44 had the highest predicted proteome coverage (33%) (**Table S5**). The latter result correlated with 16S rRNA amplicon sequencing data(19), which determined *A. muciniphila* YL44 as the most abundant bacterial species. The number of proteins detected per condition was relatively similar for each species (**Figure S4**).

From 2,860 quantified bacterial proteins, we identified 1,317 proteins whose levels significantly varied between the mice groups (ANOVA followed by THSD post-hoc analysis, FDR 5%). For four bacterial species with the largest number of detected proteins (*C. clostridioforme* YL32, *A. muciniphila* YL44, *Blautia coccoides* YL58, *Muribaculum intestinale* YL27), about 50% of the quantified proteins showed differential levels (**Table S5**), prompting in-depth evaluation of the strains’ protein profiles.

The proteomes of these bacterial species displayed an array of responses to antimicrobial-induced ecosystem perturbances (**Figure 2A**). Antibiotic or antifungal treatment appeared to have a similar effect on the proteome of *A. muciniphila* YL44 (comparison of BY_ABX vs. BY_AFX groups), while for *M. intestinale* YL27, *B. coccoides* YL58, and *C. clostridioforme* YL32, the antibiotic treatment led to more proteins with increased levels than in the antifungal groups (Table S6).

For the *A. muciniphila* YL44, *M. intestinale* YL27, and *B. coccoides* YL58, many proteins had significantly increased levels in response to treatment with either antifungal or antibiotic (B/BY compared to ABX/AFX). For *C. clostridioforme* YL32, the detected proteome showed a contrasting response to the antimicrobial treatments. For instance, compared to the fecal proteome of the B only condition, there were 2.6 and 1.3 times less proteins with elevated levels than in the BY_AFX and BY_ABX groups, respectively (**Table S6**). But when fungal species were present in the mouse gut (BY group), the effect of the antibiotic on *C. clostridioforme* YL32 appeared similar as for the other three strains (BY compared to BY_ABX), while treatment with antifungal led to a 1.3-fold increase in proteins with elevated levels in the BY group (BY compared to BY_AFX). These comparisons indicate that antimicrobial perturbance targeting bacterial or fungi cause differential responses among bacterial species that depend on species identity and presence or absence of fungi.

**Figure 2.**
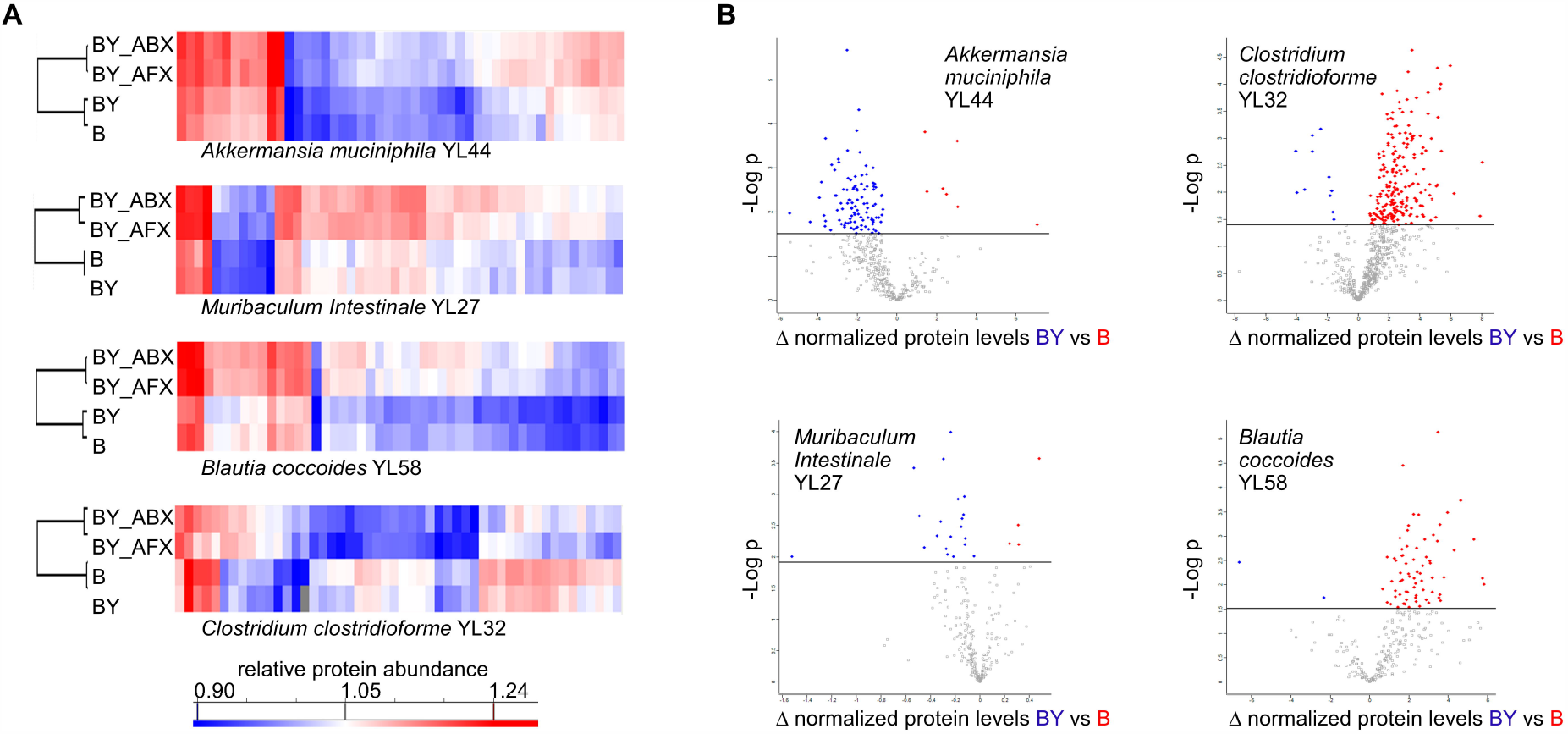
The response of four bacterial proteomes to treatment with antimicrobials and the presence of gut fungal species. A) A selection of 50 most significant differentially produced proteins for *A. muciniphila* YL44, *M. intestinale* YL27, *B. coccoides* YL58, and *C. clostridioforme* YL32 (ANOVA, FDR 0.05). B) Volcano plots representing the results of student t-test statistical comparison in protein levels between B and BY mice groups (FDR 0.05).

For the rest of the bacterial strains, low numbers of quantified proteins did not allow for in-depth investigations. Nonetheless, seven of the strains showed considerable differences in the protein profiles between the mice groups (**Figure S5)**. A general picture has emerged from this comparison, in which antimicrobial treatments targeting bacteria or fungi during the second week of the mice’s life resulted in lasting changes in the proteomes of the gut bacterial species.

### Gut fungi differentially modulate bacterial proteomes

Next, we statistically compared the bacterial protein levels between the B and BY groups to investigate the effect of fungi on the proteomes of the four strains with the highest numbers of detected proteins (**Figure 2B**). Similarly to what we observed for antimicrobials, fungal presence diversely influenced the proteomes of individual bacterial species. *A. muciniphila* YL44 and *M. intestinale* YL27 displayed increased amounts of proteins from various functional classes in the presence of fungi (**Figure S6**). We observed the opposite for *C. clostridioforme* YL32 and *B. coccoides* YL58, where the levels of 100 out of 107 and 51 out of 52 differentially produced proteins, respectively (ANOVA, THSD post-hoc, FDR 5%), decreased when fungi were present (**Table S6**). The proteomic observations for *A. muciniphila* YL44 appeared to be inversely correlated with a decrease in *A. muciniphila* abundance based on 16S rRNA sequencing in the presence of fungi (19) (**Figure S6**). For *C. clostridioforme* YL32, the trend of increased protein levels in the B group compared to the BY group also appeared to inversely correlate with the 16S-based abundances; however, these trends did not reach statistical significance in the 16S sequencing data. Similarly, we could not derive a correlation with the sequencing data for *M. intestinale* YL27 and *B. coccoides* YL58, as the strains’ relative abundance appeared similar across the mice groups. Proteomics analyses thus provided complementary information to the sequencing data and yielded a higher resolution of the ecological interactions between the gut microbial species.

### Gut bacteria and antimicrobial treatments modulate fungal proteomes

We detected 1,492 proteins for the five fungal species colonizing the gnotobiotic mice, and 39% of these proteins were assigned LFQ levels. The lower numbers of proteins detected and quantified, as compared to bacteria, were a result of a lower abundance of fungi in the gut (19). Most of the identified fungal proteins were present in the Y group representing the mice colonized exclusively with fungi. We previously showed that the Y group harboured higher fungal concentration than co-colonized mice (19), confirming that a bacterial suppression of the fungal colonization occurs in the host gut (36). Of the five fungal species, *C. glabrata* had the highest number of identified proteins, which also dominated the Y group (**Figure 3A**). However, antibiotic treatment (BY_ABX) led to higher detection of proteins derived from *Pichia kudriavzevii*/*C. krusei*, followed by *C. albicans and C. parapsilosis*. The total number of differentially abundant proteins between the mice groups and their distribution among the fungal species were rather similar for the following pairs of mice groups: Y - BY_ABX and BY - BY_AFX (**Figure 3B)**. When we expressed the number of differentially produced fungal proteins as a percentage of each strains detected proteins, we noticed that *C. glabrata* proteome showed a stronger response to the antibiotic treatment (BY_ABX), as compared to Y, BY and BY_AFX.

**Figure 3.**
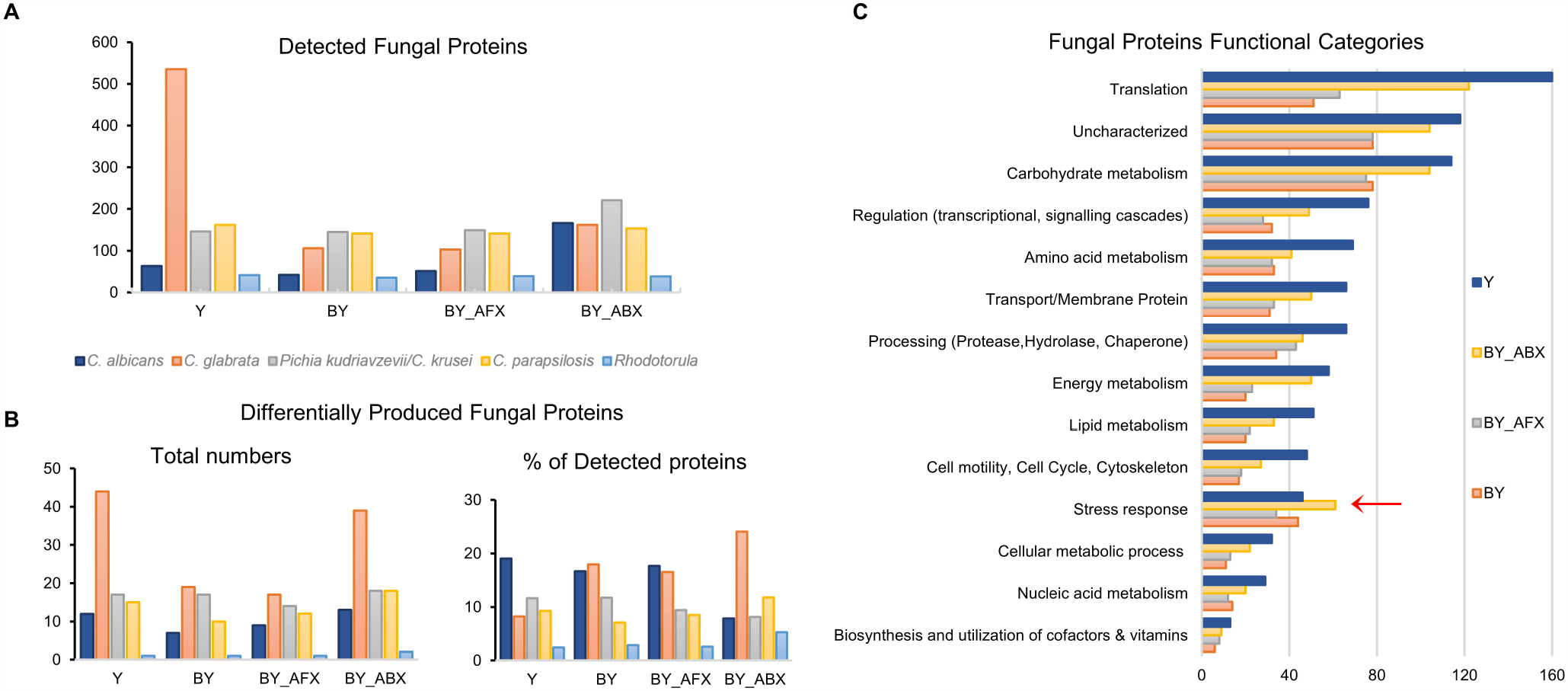
Response of fungal proteomes to antimicrobials and presence of bacteria. A) Distribution of detected fungal proteins across four mice group (n=1,492). B) Distribution of fungal proteins with significantly changed levels across four mice groups (n=99, ANOVA, FDR 5%). C) Distribution of the main functional classes of detected fungal proteins (n=1,492). Proteins functional annotation was downloaded from the UniProtKB database (29) and compared to annotations obtained using the DAVID (30) and STRING-db (31) tools.

The presence of bacteria and early-life antimicrobials had significant effects on the fungal species protein profiles. Arrangement into categories using functional annotations based on UniProtKB (29) and other databases (30, 31), showed an increased number of fungal proteins related to stress responses in the antibiotic condition (**Figure 3C**), suggesting that the fungal species are either negatively affected by the changes in the bacterial microbiome or by the antibiotic treatment itself. Interestingly, many of the identified fungal proteins were previously reported to be excreted via extracellular vesicles (37, 38) and to have immunogenic properties (39, 40) (**Table S7**). Among these proteins were enzymes of the glycolytic pathway (*e*.*g*., enolase, glyceraldehyde-3-phosphate dehydrogenase, fructose-bisphosphate aldolase, and phosphoglycerate mutase) and molecular chaperones linked to stress response (heat shock proteins). These proteins were significantly elevated in the fungal group suggesting that they are critical cytosolic proteins abundantly produced by fungal cells living in the mouse gut.

### Fungal and bacterial colonization induces distinct and persistent changes in the host fecal proteome

We identified 674 mouse proteins as part of the mice fecal metaproteome. From proteins quantified by LFQ (405), 71% displayed differential abundance between the groups (ANOVA followed by THSD post-hoc analysis, FDR 5%, **Table S8**). PCA score plot based on the quantified proteins showed four clusters (**Figure 4A**): the first contained co-colonized mouse groups treated with antimicrobials (BY_ABX/AFX), and the second consisted of the B and BY groups. The third and fourth clusters included the Y and GF groups, and these were considerably more dissimilar from the first two clusters. Host proteins in the Y group were more clearly separated from the GF mice, in contrast to clustering when also microbial proteins were considered (**Figure 1C**), revealing that the proteome response to fungal colonization may be more pronounced in the host proteome than in microbial cells. The GF group displayed the highest number of proteins associated with lipid metabolism, regulation, and molecular processing, while the BY and BY_AFX groups had the highest numbers of immune proteins (**Figure 4B**). In the BY_ABX group, we identified an increased number of stress response proteins. Moreover, the Y and GF groups displayed an increased number of proteins associated with energy metabolism (*e*.*g*., mitochondrial proteins).

**Figure 4.**
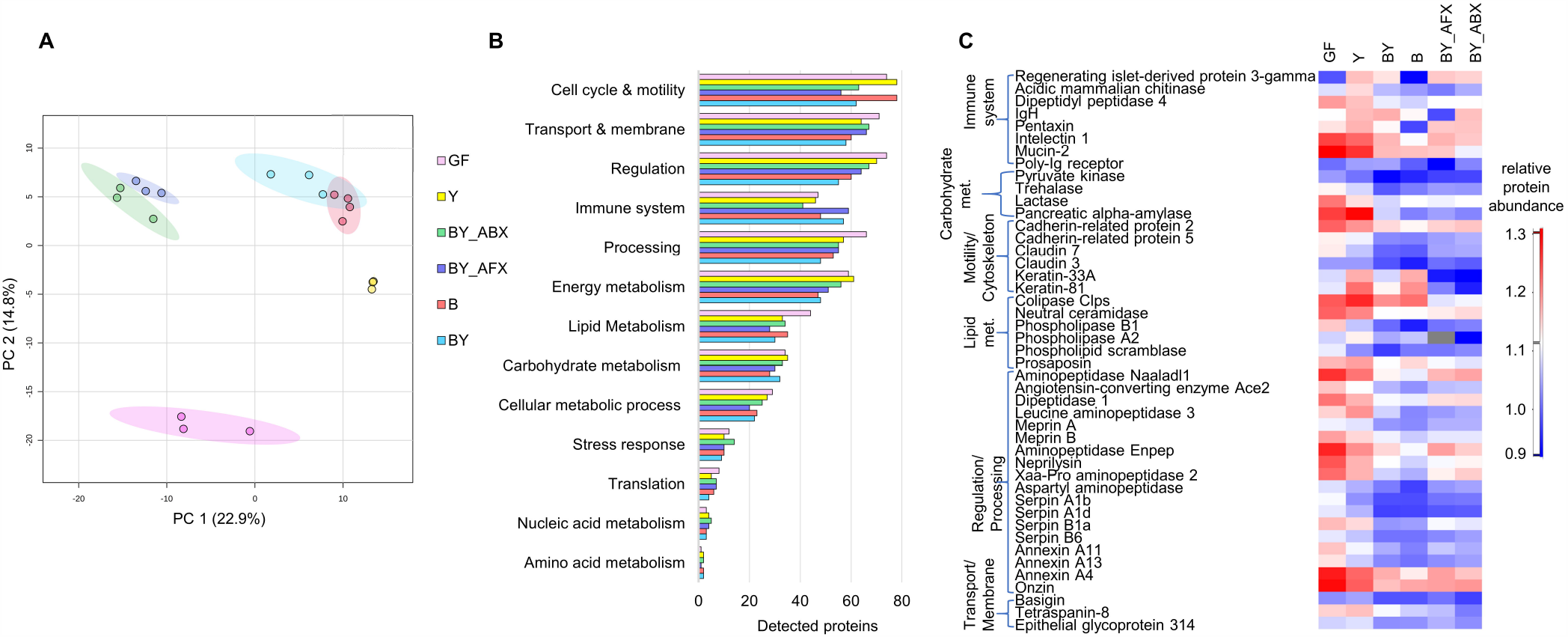
The response of host fecal proteome to microbial colonization and antimicrobial treatment. A) PCA score plot based on 405 quantified mouse proteins. X- and Y-axis show the first and second principal components, accounting for 22.9% and 14.8% of the total variation, respectively. B) Functional classes of 651 detected mouse proteins, with the protein annotations derived from the UniProtKB database (29), and compared to those obtained by the DAVID (30) and STRING-db (31) tools. C) Selection of differentially produced mouse proteins extracted from feces. Abbreviations of the mice treatment groups: B, bacteria; BY, bacteria + fungi; BY_ABX/AFX, bacteria + fungi and antibiotic or antifungal treatment; GF, germ-free; Y, fungi/yeast.

Proteins whose levels significantly varied between the mouse groups belong to the following functional classes: processing proteins such as proteases, hydrolases, and chaperones (14%), proteins linked to cell cycle regulation, motility, and cytoskeleton, including components of the tight junctions (14%), transport and membrane proteins including epithelial cell surface receptors (13%), proteins of energy and lipid metabolism (12% and 9%, respectively), and gut immune and barrier factors (9%). To tease apart the mouse proteome response to different microbes, we statistically and functionally compared pairs of treatment groups containing either fungi or bacteria (**Figure S7**). The introduction of the 12-species bacterial consortium led to a striking decrease in the mouse protein levels, and this was valid both for germ-free mice (GF vs. B) and mice colonized with fungi (Y vs. BY). Although gut fungi did not have the same significant effect on the mouse proteome as bacteria, we detected a 5-fold decrease in the number of proteins whose levels increased in the Y group compared to GF (GF vs. Y),. The presence of fungi and bacteria resulted in a 2-fold increase compared to bacteria alone (BY vs. B), particularly of proteins associated with the immune system, energy metabolism, and cellular processing.

Because the mouse protein levels were strongly dependent on the gut microbial consortium, we gave special attention to the individual functional classes of differentially produced proteins. Levels of 27 immune proteins were the highest in GF mice, followed by those colonized only with fungi (**Figure 4C**). Two immune proteins were elevated in the presence of fungi: acid mammalian chitinase (Chia), implicated in the defense response against fungi, and regenerating islet-derived protein 3-gamma (Reg3g), a bactericidal C-type lectin reported to act exclusively against Gram-positive bacteria. Thus, our results suggest that Reg3g is also involved in the intestinal cells’ response to fungal colonization.

Bacterial presence resulted in a reduction in the mucosal Pentaxin (Mtx2) levels, a secreted protein involved in complement activation, Intelectin 1 (Itln1), a receptor that binds microbial glycans, and Dipeptidyl peptidase 4 (Dpp4), cell surface receptor and dipeptidyl protease involved in T-cell activation. All these proteins had the lowest amounts in the B group. Levels of other immune proteins, such as Mucin 2 (Muc2), poly-Ig receptor (Pigr), and immunoglobulin heavy chain (IgH), were altered by the shifts in microbial composition caused by antimicrobial treatments.

The absence of microbes resulted in elevated levels of mouse proteins linked to carbohydrate metabolism both at the cellular [*e*.*g*., Pyruvate Kinase (Pkr)] and host level [glycosidases such as Pancreatic Alpha-Amylase (Amy2), Lactase (Lct), and Trehalase (Treh)]. Amy2, a glycosidase that hydrolyses alpha-linked polysaccharides such as starch and glycogen, had the highest levels in the Y group, suggestive of increased Amy2 production response to either fungal polysaccharides or fungal metabolism on dietary sugars. Fungi also elicited a strong effect on host proteins associated with lipid metabolism. The highest levels were identified in the Y group for the host enzyme Colipase (Clps), a cofactor of pancreatic lipase facilitating lipids digestion, and Phospholipase A2 (Pla2g1b), a secreted protein involved in lipid degradation and innate immune mucosal response. In contrast, membrane-associated Phospholipase B1 displayed the highest and lowest levels in the GF and B group, respectively. Closely connected to the immune system are phospho- and sphingolipid metabolism enzymes, which often have regulatory properties towards immune cells. Of those, we identified Neutral Ceramidase (Asah2), a membrane protein hydrolysing sphingolipid ceramides into sphingosines and free fatty acids, whose relative levels were the lowest in BY and B groups. Phospholipid Scramblase (Plscr1), an intracellular protein mediating migration of phospholipids, and Prosaposin (Psap), a secreted protein involved in sphingolipid metabolism, both showed the lowest levels in the B group.

Cytoskeletal and cell adhesion proteins were also significantly increased in the GF and Y mice. These included cadherin-related proteins 2 and 5 (Cdhr2 and Cdhr5), which regulate microvilli length, and Claudins 3 and 7 (Cldn3 and Cldn7) controlling tight junction-specific obliteration of the intercellular space. Notably, Keratin-33A and −81 (Krt33a and Krt81), structural proteins that form the cytoskeleton’s intermediate filaments in epithelial cells, displayed the highest levels in the Y group, followed by the B group and had the lowest levels in the BY groups treated with antimicrobials. Adhesion proteins with regulatory properties and links to the immune system included Tetraspanin-8 (Tspan 8), Epithelial Glycoprotein 314 (Epcam), and Basigin (Bsg), a cell surface receptor. All three proteins showed increased levels in the GF and Y groups.

One of the most prevalent functional classes among the differentially produced proteins were proteases, which had the highest levels in the GF and Y groups, and the lowest in the BY and B groups. These included aminopeptidase Naaladl1, Dipeptidase (Dpep1), Cytosol/Leucine aminopeptidase 3 (Lap3), Xaa-Pro aminopeptidase 2 (Xpnpep2), Aspartyl aminopeptidase (Dnpep), Aminopeptidase (Enpep), Neprilysin (Mme), and Angiotensin-converting enzyme (Ace2). Interestingly, Ace2 is a carboxypeptidase with multiple regulatory functions, including gap junction assembly. Other proteases with regulatory properties included membrane-bound metalloproteases Meprin A (Mep1a) and B (Mep1b), implicated in the inflammatory response. Alongside proteases, protease inhibitors followed the same quantitative pattern as described above. These included Serpins, serine protease inhibitors that negatively regulate endopeptidase activity in response to cytokines (Serpinb1a, Serpina1d), innate immune response, inflammation, and cellular homeostasis (Serpinb1a, Serpinb6).

Numerous regulatory proteins were present among the proteins with increased levels in the GF and Y group, such as the Annexin family of Ca2+-regulated phospholipid-binding and membrane-binding proteins (Anxa4, Anxa11, Anxa13) and nuclear proteins such as Onzin (Plac8) suggested to regulate immune responses. Overall, the metaproteomic analyses documented an extensive impact of microbial colonization in a controlled early life gut microbiome model, underlining the intertwined functional development of the host and its gut microbiome, and revealing new features of the host response to fungal colonization.

### Fungal colonization drives alterations in the mouse jejunal tissue proteome

Given the exciting findings from the mice fecal metaproteomes, we decided to investigate more closely the host proteome. We selected the small intestine as our site of interest because it contains the body’s largest immune organ (the gut-associated lymphoid tissue) and is an important site of the host-microbe interactions (41). We used a tandem mass tag labelling approach (TMT 6-plex) to gain a higher sensitivity for low abundant proteins (**Figure 5A**). From the TMT 6-plex experiment, we identified 10,201 peptides (**Table S9)** matched to 1,514 mouse proteins (**Table S10)**, and the levels of 45 were significantly altered between the mice groups (ANOVA, THSD post-hoc analysis, FDR 5%, **Table S11**). PCA score plot revealed different grouping results than for the mouse fecal proteomes, as we observed four distinct clusters (**Figure 5B**). Functionally, the 45 significantly changed proteins were mainly associated with cellular metabolism and regulation of cellular processes (**Figure 5C**).

**Figure 5.**
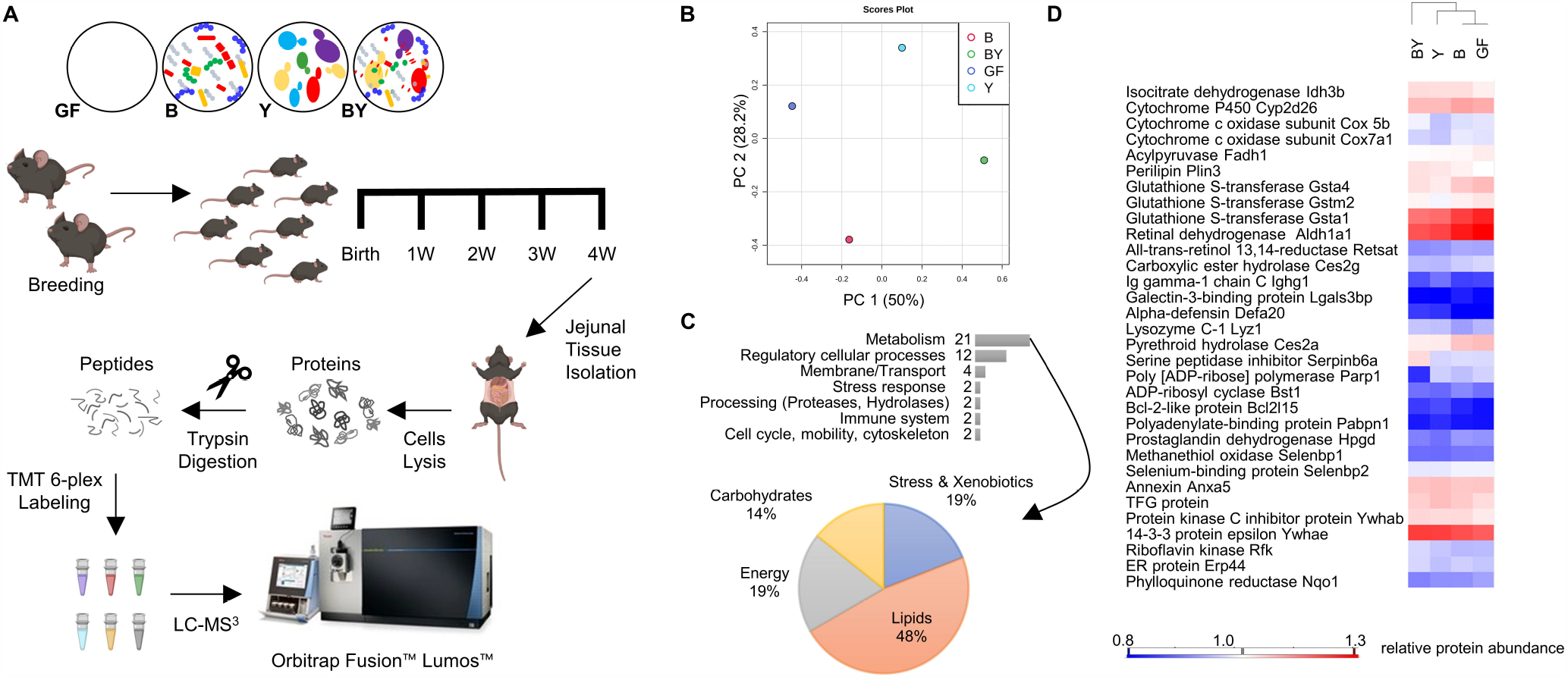
Response of mouse jejunal proteomes to microbial colonization. A) Experimental design of Tandem Mass Tag (TMT) labelling experiment. B) PCA score plot based on 1,377 mouse proteins quantified in a minimum of 4 biological replicates (replicates mean shown). X- and Y-axis show the first and second principal components, accounting for 50% and 28.2% of the total variation, respectively. C) Functional classes of 45 differentially produced mouse jejunal proteins (ANOVA followed by THSD, FDR 5%). D) Selection of differentially produced mouse proteins extracted from jejunal tissue. Abbreviations of the mice treatment groups: B, bacteria; BY, bacteria + fungi; GF, germ-free; Y, fungi/yeast

Compared to the fecal mouse proteome, proteome changes of the jejunal tissue related to microbial colonization were more subtle (**Figure 5D**). Among immune proteins whose levels increased with the presence of fungi were Alpha-defensin 2 (Def20) and Ig gamma-1 chain C region secreted form (Ighg1), while Galectin-3-binding protein (Lgals3bp) had the highest levels in the bacteria group. From 10 proteins functionally connected to lipid metabolism, two Glutathione S-transferase enzymes (Gstm2 and Gsta4) were significantly decreased in the fungal group, and yet another Glutathione S-transferase (Gsta2) showed differential levels across the mice groups. We also detected two proteins of retinol metabolism (Retsat and Aldh1a1), which play a key role in mucosal immune responses. Fungal colonization further appeared to impact the intestinal cells’ energy metabolism, as several mitochondrial proteins had decreased levels in the fungi group, most notably, two subunits of Cytochrome C oxidase (Cox5b and Cox7a1).

Some of the differentially produced proteins have functional links to NF-κB pathway, such as the apoptotic marker Poly (ADP-ribose) polymerase (Parp1). In some contexts, particularly in response to cellular stress, stimulation of the NF-κB pathway promotes apoptosis (42). Selenium acts as a key element that controls NF-κB activation and the half-life of its inhibitor IκBα, and we detected two selenium-binding proteins (Selenbp1 and Selenbp2). Other proteins associated with NF-κB signalling included Trafficking from ER To Golgi Regulator (Tfg), Bcl-2-like protein 15 (Bcl2l15), Prostaglandin dehydrogenase 1 (Hpgd), Protein kinase C inhibitor 1 (Ywhab), and Annexin A5 (Anxa5). Overall, the TMT 6-plex analysis of the jejunal tissue provided further insight on host cellular pathways impacted by defined microbial colonization and confirmed that gut fungi elicit differential effects compared to bacteria.

## DISCUSSION

In this work, we used a controlled environment of gnotobiotic mice colonized with defined microbial consortia to precisely describe changes in the metaproteome profiles associated with alterations in the gut microbiome. By reducing the gut microbiome’s complexity to 12-bacterial species, five fungal species, or the combination of the 17 microbes, the gnotobiotic model allowed us to isolate specific contributions and cross-kingdom interactions between the gut-associated bacteria and fungi, as well as the host response to microbial colonization.

Our previous work determined the impact of gut fungi on gut microbiome community structure and host immune development (19). Here, we strengthened these results using a metaproteomic approach that provided an independent functional measure of the microbiome-host crosstalk. Similarly to other multi-omics studies (43, 44), we found discrepancies between the quantitative proteome profiles and the DNA-based relative abundance of individual species. These differences likely stem from complex regulatory networks along the gene to protein expression path, which differ between microbial kingdoms. The latter was recently documented in a study showing dynamic ratios between protein and RNA levels that depended on the type of microbial population (45).

For the fungal species, a meaningful comparison between the protein and DNA data was hindered by the absence of genome-based protein databases that would guide the raw MS data searches. However, corresponding with our earlier results (19), the proteomic data showed that fungi grew in higher concentration in bacteria’s absence. This finding is in line with the interkingdom competition and antagonism between bacteria and fungi in other ecosystems, such as the rhizosphere and soil (46), and in mammalian hosts, where commensal bacteria limit fungal colonization via activation of innate mucosal immunity (36) and directly by producing inhibitory metabolites such as short-chain fatty acids (47).

Despite the lack of a linear relationship between the DNA and protein data, our earlier results from the 16S rRNA sequencing aided the interpretation of the bacterial species proteome responses, and the combined analyses revealed antagonistic and synergistic relationships between the bacterial and fungal species. For example, we identified *Lactobacillus reuteri* as a potent responder to the fungal presence from the sequencing data, as the bacterium abundance was significantly reduced in the BY group. Only a few *L. reuteri* proteins were quantified, and those displayed lower amounts in the absence of fungi, conversely to the DNA sequencing results (**Figure 6A**). Some of the most abundant bacterial proteins were detected in the B-only group (ribosomal protein L31, enolase, and translational factor Tu) along with three dehydrogenases, whose increased production often indicates cellular stress. For another low abundant bacterial species, *Clostridium innocuum*, we observed a similar reciprocal relation between the proteomic and sequencing data (**Figure 6B**). Taken together with the observed variations in the proteome response of the four most abundant bacterial species (**Figure 2**), the proteomic data is suggestive that the cells are negatively affected by changes in the gut environment related to the fungal presence (*L. reuteri, A. muciniphila, M. intestinale*) or absence (*C. innocuum, B. coccoides, C. clostridioforme*). Similar results were found in samples from mice treated with antimicrobials, which affected levels of numerous proteins and often led to their elevated production (**Figure 2 and S5**). Such proteome response has been documented for bacterial cells treated with antibiotics (48), where the cells optimize their proteins’ production to deal with the adverse environmental factor.

**Figure 6.**
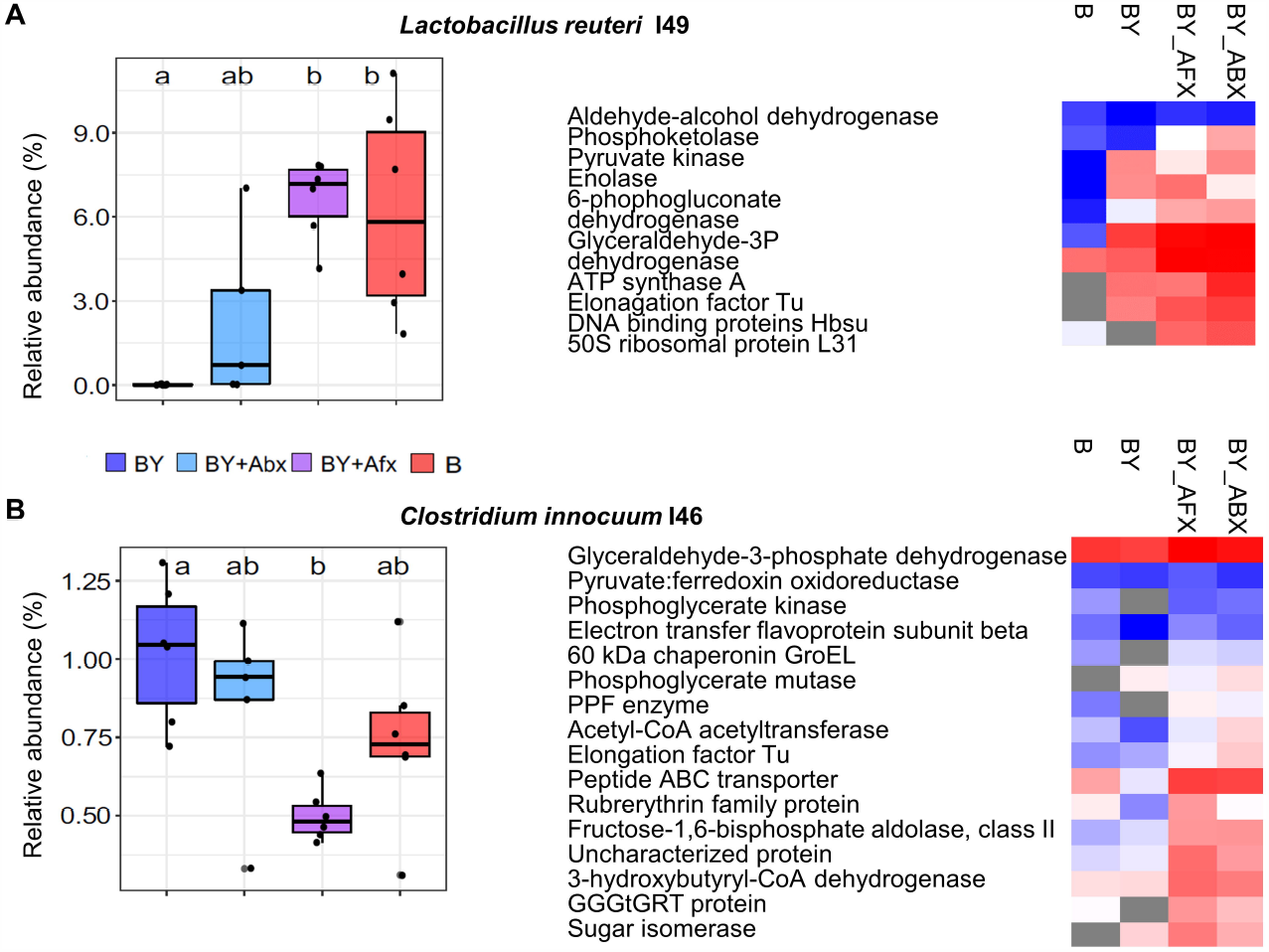
Detection of stress response in bacterial proteomes. The relative abundance based on 16S rRNA sequencing and differentially produce proteins of *Lactobacillus reuteri* (A) and *Clostridium innocuum* (B). Proteins quantified in a minimum of two replicates are shown; grey fields in heatmaps indicate the protein amount was below the quantification limit. Colour denotes microbial colonization (red B - bacteria, blue BY – bacteria + yeast, cyan blue BY_Abx – BY + antibiotic, purple BY_Afx - BY + antifungal).

One of this study’s most striking findings was a similar impact of the antibacterial and antifungal treatments on bacterial and fungal species’ proteomes. Inhibitory effects of antifungal drugs on the growth of commensal bacteria are poorly described (49), but it may be that the antifungal Fluconazole inhibits intestinal bacteria to some extent via direct or indirect mechanisms, similarly to other non-antibiotic drugs (50). The antibiotic and antifungal treatments also had a strong effect on the quantities and types of fungal proteins being produced, revealing how these antimicrobial drugs impact the entire community, with similar functional consequences. Ecosystem perturbances caused by either antibiotics or antifungal drugs may impact the gut microbiome keystone species, causing a widespread effect on microbial networks and foodwebs (51).

Host proteins present in the intestine are essential for maintaining a mutualistic relationship with the microbes on the mucosal interface and serve as reporters on the host-microbe interactions. Host fecal proteome has been reported to exhibit signatures specific to colonization states (52). However, to our knowledge, protein response to exclusive fungal colonization has not been characterized before. For the first time, we described at the proteome level the cellular pathways and their protein components involved in the interactions between the mammalian host and gut fungal species. Fungal colonization resulted in changes in host proteins functional in innate immunity as well as metabolism (**Figure 4**), suggesting specific roles of gut fungi on host systems during early developmental stages. Further research aimed at investigating these roles has great potential for novel discovery, given that the majority of host-microbiome interactions described to date have been limited to bacteria.

In the host jejunal proteome, most of the differentially produced proteins had metabolic functions (**Figure 5**). The quantitative profiles of proteins with significantly changed levels showed closer clustering of the GF and B groups (**Figure 5D**). These results likely reflect the host control of the number of bacterial cells in the small intestine, where they would otherwise compete for easily digestible nutrients. Alternatively, the more subtle differences in jejunal proteomes across microbial colonization conditions, as compared to profound changes identified in fecal metaproteomes, may also reflect host homeostatic pressures present in intestinal tissues. Nonetheless, these findings indicate that fungal colonization influences host jejunal proteome in a distinct way from bacteria.

The identification of multiple fungal proteins hypothesized to be abundant components of the extracellular vesicles suggests that fungi influence the host through direct cell-to-cell contact. Possible mechanisms may be akin to recently described interactions between segmented filamentous bacteria and mice intestinal epithelial cells (53). These bacteria protruded into the epithelial cells and used adhesion-triggered endocytosis to transfer antigens into intestinal epithelial cells and modulate host T cell homeostasis. Similarly, epithelial internalization of fungal hyphae, a morphology into which fungal cells transition to strengthen their adherence to epithelial cells (54), can deploy fungal extracellular vesicles with host cell-modulatory properties. Fungal cells produce a diverse array of biologically active compounds (55, 56), but the understanding of the mycobiome contribution to immunomodulatory substances present in the gut remains rudimentary. Our work sets the stage for future studies that will explore the functional mechanisms associated with gut fungal colonization.

Limitations of this work include the lack of genome references for the fungal strains colonizing the mice. Further efforts to carry out whole genome sequencing of fungal species and strains commonly associated with mammalian gut will reveal a better-resolved analysis of the functional role of these understudied microbes to their microbial ecosystems and their impact on the host. Also, fecal samples are a proxy for the colon microbiome. Although technically challenging, sampling along the gastrointestinal tract will reveal additional insights on microbial colonization of the small and large intestine. Finally, label-free quantification is a common strategy adopted in metaproteomic studies but has low sensitivity to more scarce proteins as well as limitations in quantification accuracy. The application of metabolic labeling for improved peptide quantification (57) holds the potential to improve the accuracy of quantitative metaproteomics, and promote the application of proteomics for functional studies of intestinal microbiomes.

## ACKNOWLEDGEMENTS

We thank Erik van Tilburg Bernardes and Isabelle Laforest-Lapointe for providing the mice fecal samples, and Ali Javed for a contribution to the fungal proteomes’ analyses, all at the time at the Arrieta lab. We also thank Daniel Young from the Dufour lab for assistance with the proteomic sample preparations. We acknowledge the Southern Alberta Mass Spectrometry Facility and Laurent Brechenmacher for the service and support for the LC-MS/MS experiments. This work was funded by The Research Council of Norway Grant No. 274296 (V.K.P). This work was also supported by the Cumming School of Medicine, the Alberta Children Hospital Research Institute, the Snyder Institute of Chronic Diseases, the Canadian Institutes for Health Research, the Sick Kids Foundation, the and W. Garfield Weston Foundation (M.C.A). A.D. was supported by an NSERC Discovery Grant (DGECR-2019-00112). The funders had no role in the study design, data collection and analysis, decision to publish, or preparation of the manuscript.

