## Supplementary Information for "Metaproteomic Profiling of Fungal Gut Colonization in Gnotobiotic Mice"

**TABLE OF CONTENT:**

**Figure S1** TMT labels carried by each sample and the mixing design
**Figure S2** Summary of the TMT MS3 acquisition method
**Figure S3** Summary of the LFQ MS/MS acquisition method
**Figure S4** Detected bacterial proteins per strain per condition
**Figure S5** Response of bacterial proteomes to antimicrobials and the presence of fungi (seven strains with a low number of quantified proteins
**Figure S6** Number and functional classes of differentially detected bacterial proteins in either B or BY mice groups for four bacterial species.
**Figure S7.** Response of mouse fecal proteome to the presence of microbial consortiums.
**Table S6** Number of proteins with significantly increased levels for selected bacterial strains and between different mice groups

**Tables in xlxs file Supplementary Tables**

**Table S1** Microbial species used for gnotobiotic mice colonization and corresponding protein databases used in MS data search.
**Table S2** Pearson coefficients describing correlation of proteins identification and quantification across replicates.

**Table S3** Peptide sequences detected at 1% FDR (xlsx file Table_S3_Peptides_Fecal_Metaproteomics)

**Table S4** Fecal Proteins detected at 1% FDR, including LFQ intensities

**Table S5** Overview of detected proteomes

**Table S7** Fungal proteins reported as part of extracellular vesicles

**Table S8** Fecal mouse proteins with significantly differential abundance as determined by ANOVA (FDR 0.05)

**Table S9** Peptide sequences detected at 1% FDR (Jejunal Tissue)

**Table S10** Proteins from jejunal tissue detected at 1% FDR

**Table S11** Jejunal mouse proteins with significantly differential abundance as determined by ANOVA (FDR 0.05) and posthoc THSD analysis

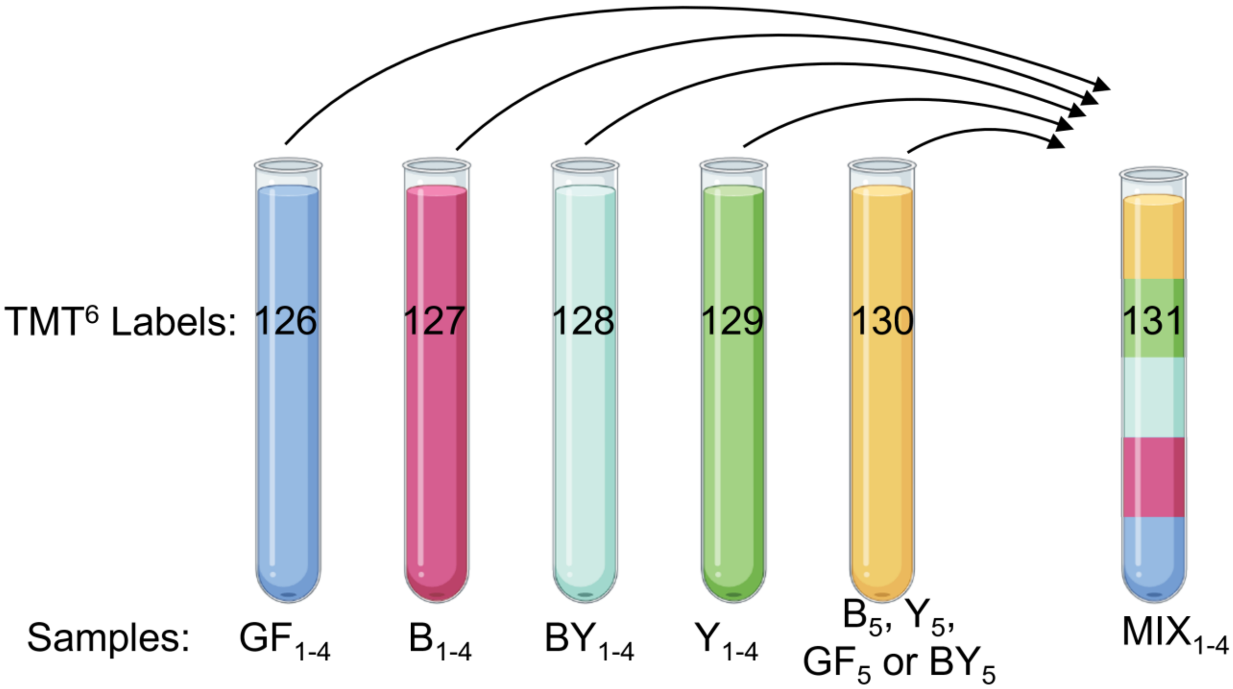


**Figure S1** TMT labels carried by each sample and the mixing design. Four TMTsixplex™ Isobaric Mass Tagging Kit were used for labelling of 20 samples and 4 pooled controls. Samples originated from jejunum of mice that were either left germ-free (GF), colonized with bacterial (B), fungi (F) or both (BY). A unique reporter mass (126-131 Da) of the TMT6 isobaric labels is described on each tube.

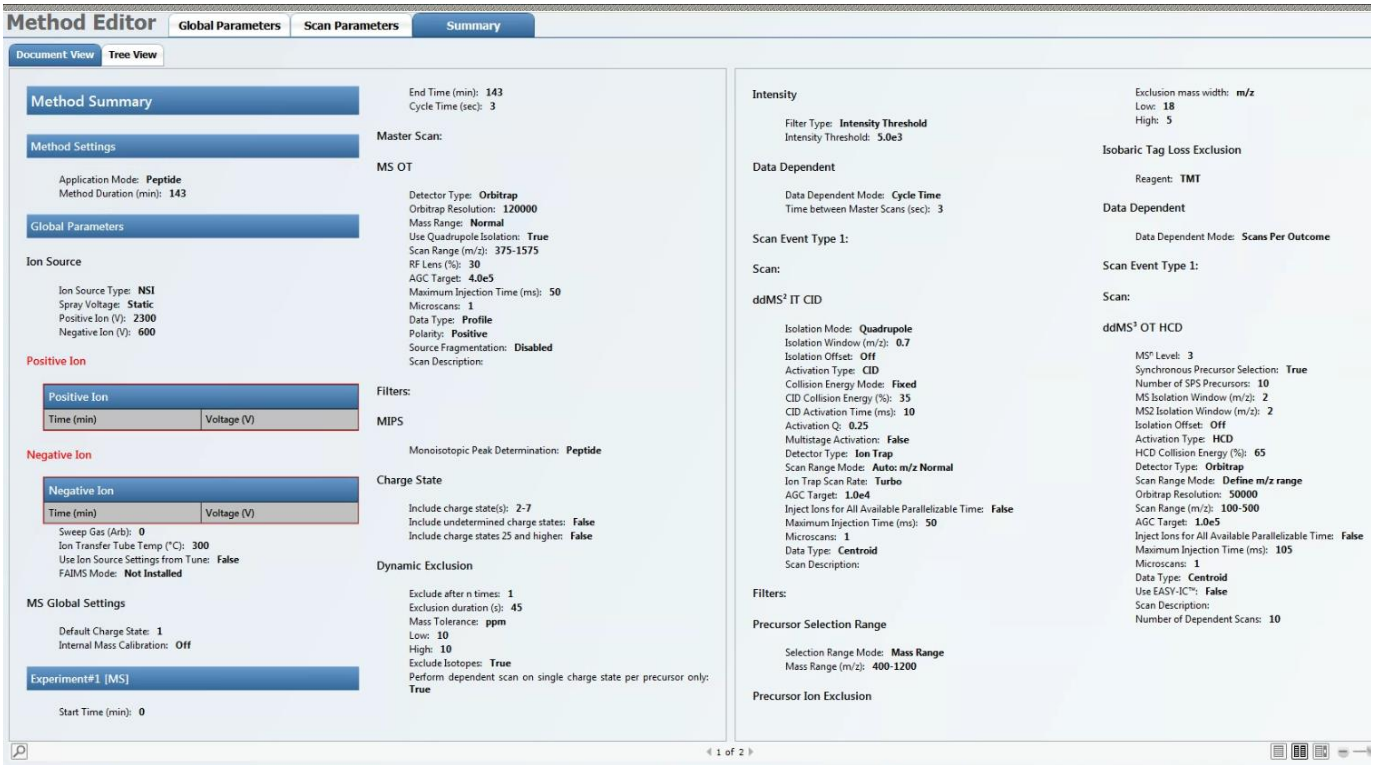


Figure S2 Summary of the TMT MS3 acquisition method

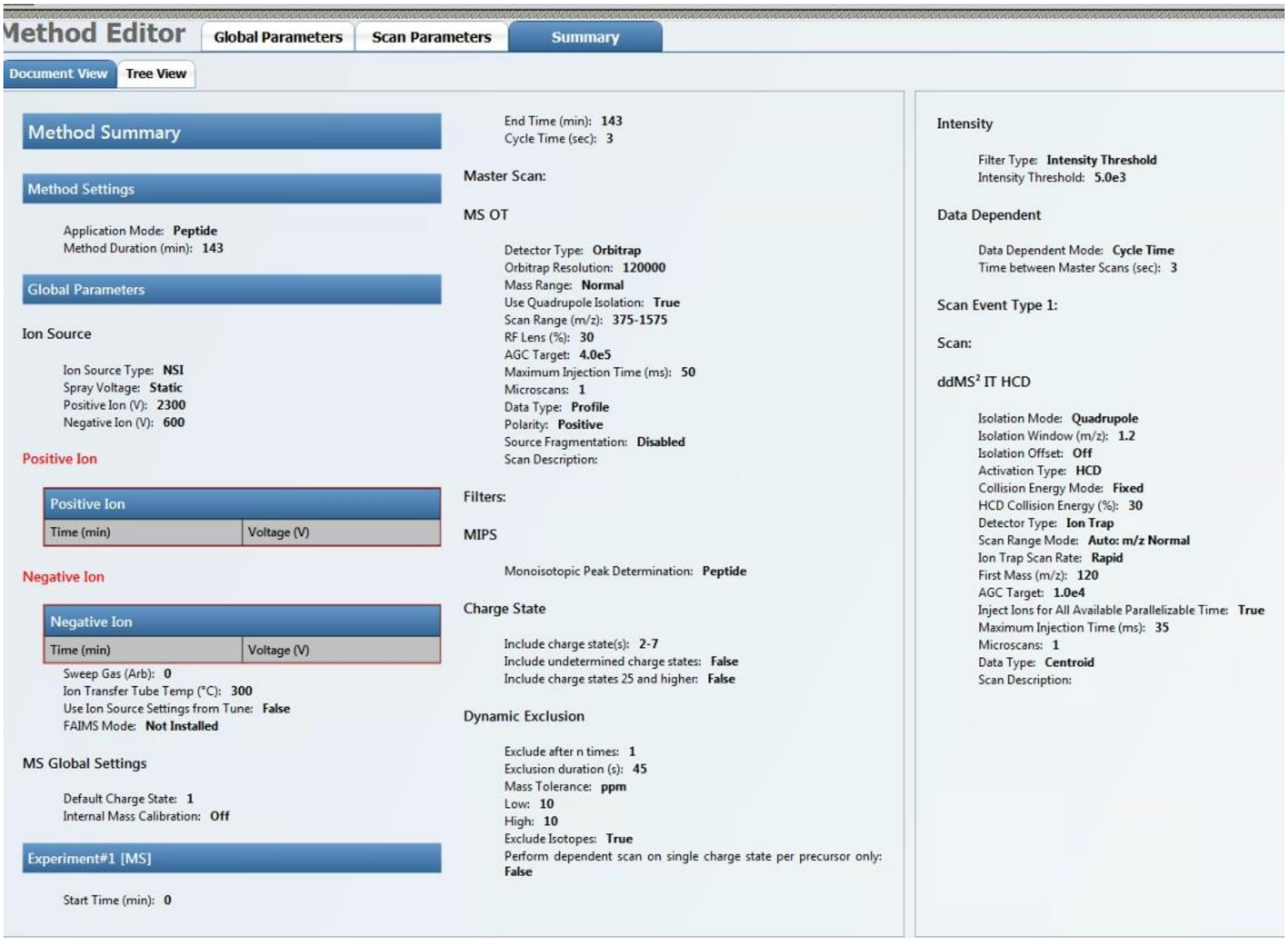


**Figure S3** Summary of the LFQ MS/MS acquisition method

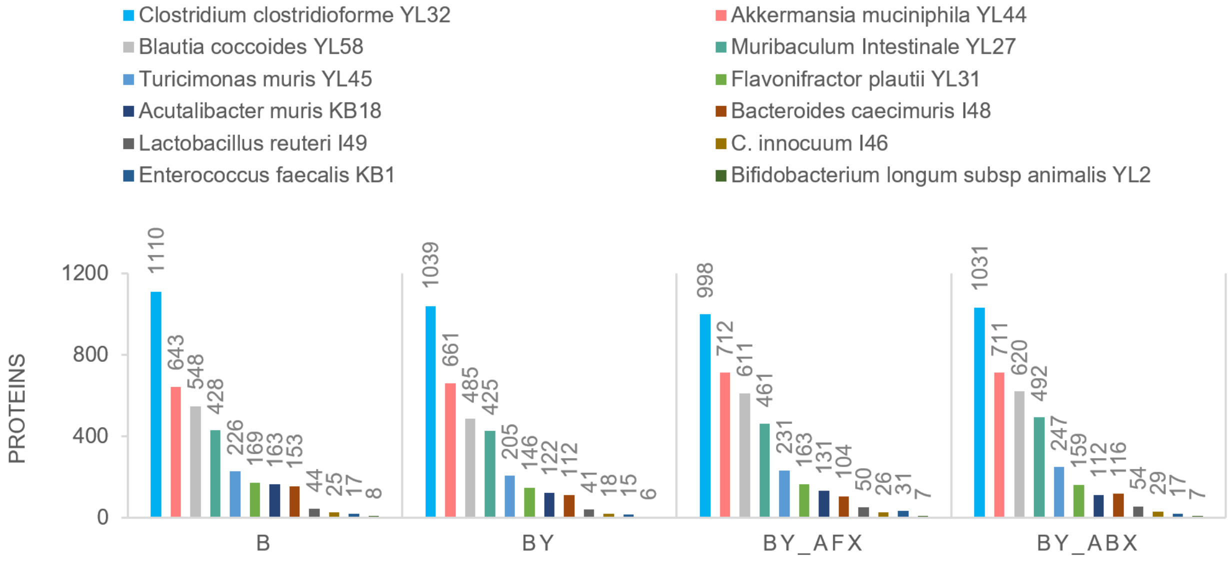


**Figure S4**. Detected bacterial proteins per strain per condition. Abbreviations of the mice treatment groups: B, bacteria; BY, bacteria-yeast; BY_ABX/AFX, bacteria-yeast, and antibiotic or antifungal treatment.

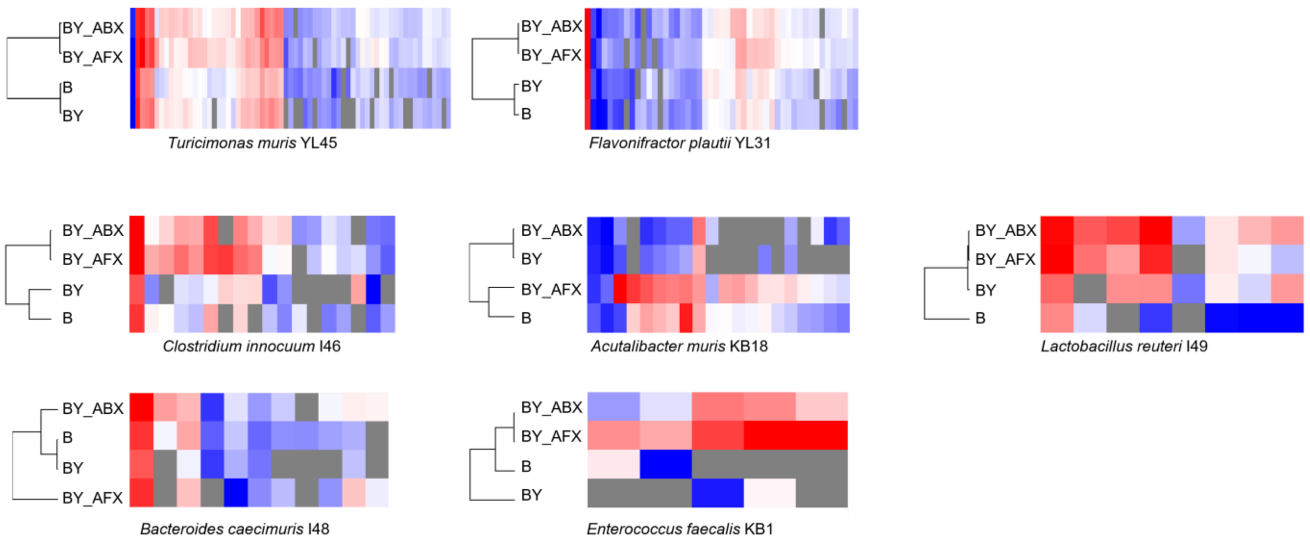


**Figure S5** Bacterial proteomes response to antimicrobials and the presence of fungi – seven strains with a low number of quantified proteins. Abbreviations of the mice treatment groups: B, bacteria; BY, bacteria-yeast; BY_ABX/AFX, bacteria-yeast, and antibiotic or antifungal treatment.

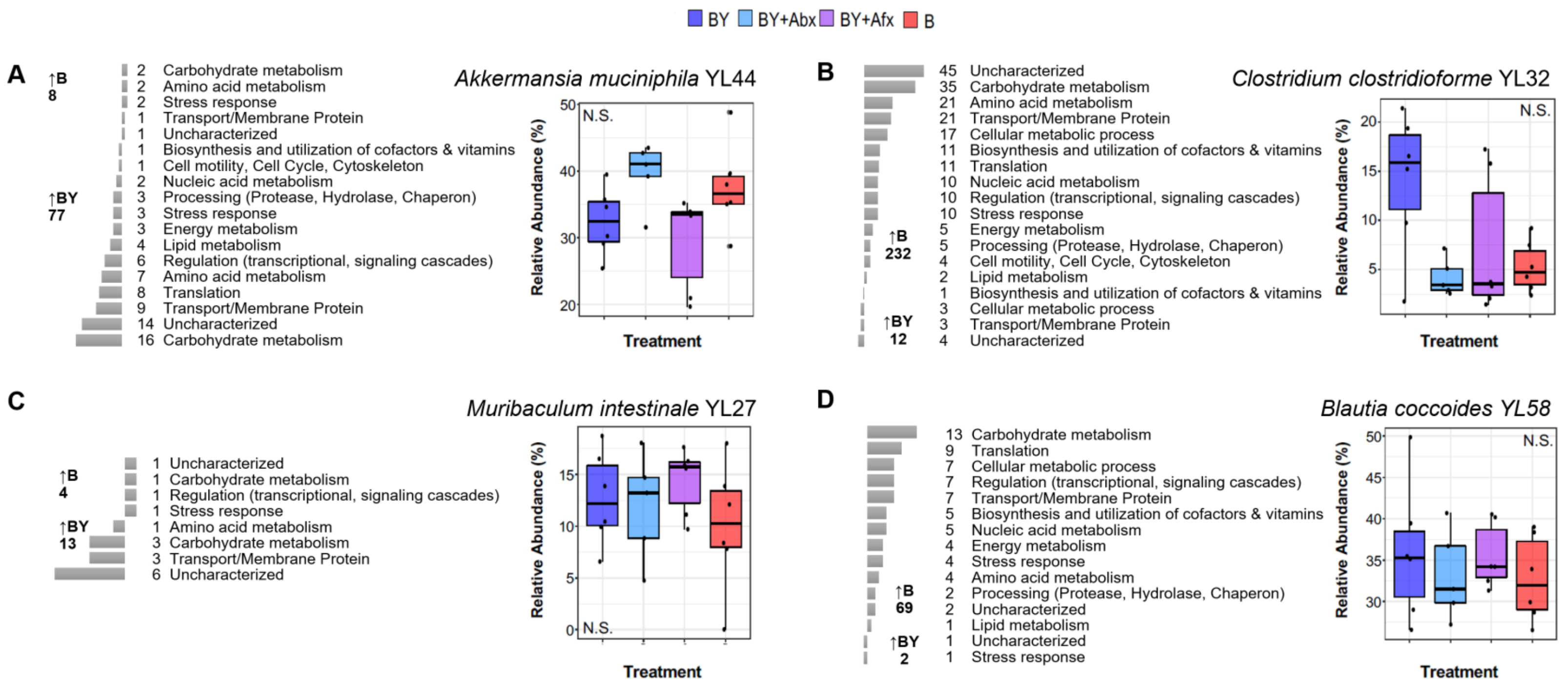


**Figure S6.** Number and functional classes of differentially detected bacterial proteins in either B or BY mice groups for A) *A. muciniphila* YL44, B) *C. clostridioforme* YL32, C) *M. intestinale* YL27, and D) *B. coccoides* YL58. Protein functional annotation was downloaded from the UniProtKB database[29] and compared to annotations obtained using the DAVID[30] and STRING-db[31] tools. The relative abundance of the strains based on 16s rRNA sequencing data is shown on each panel's right. Number of proteins in each functional category is indicated next to the bar plots (scale not comparable between A-D). N.S. – not significant.

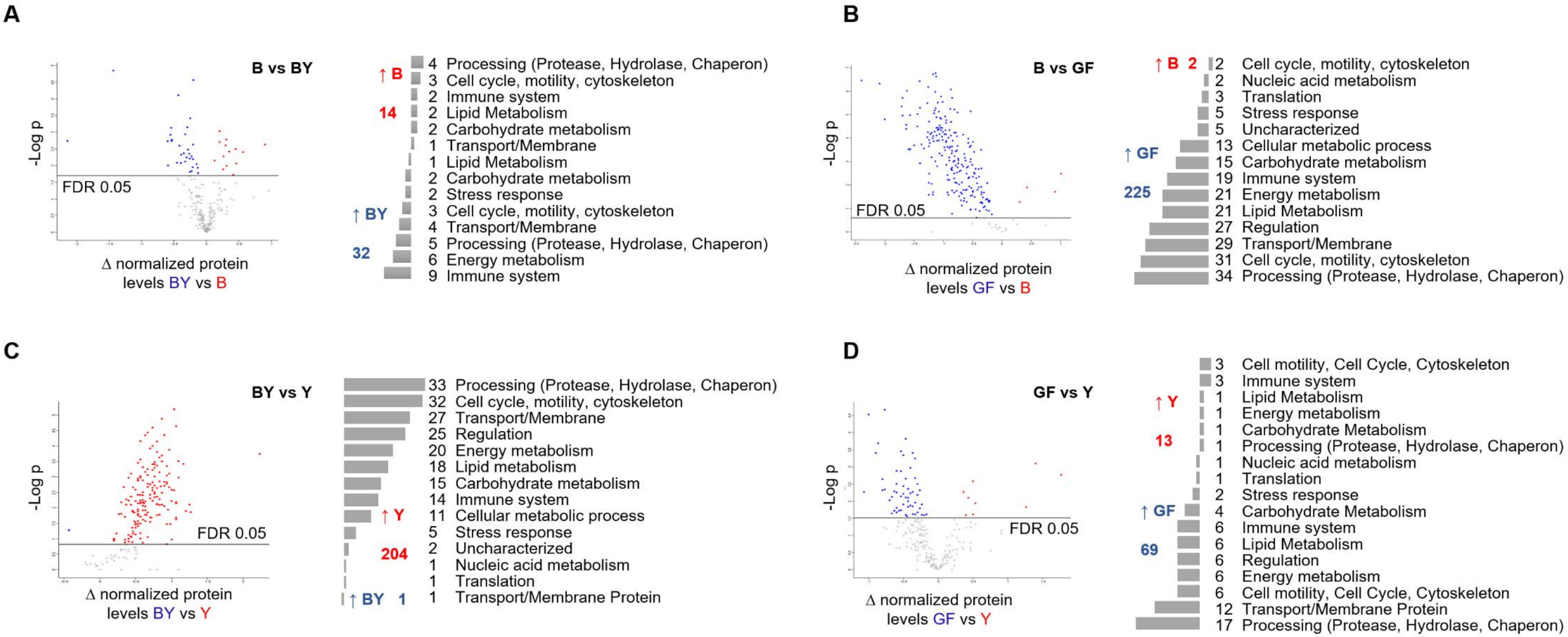


**Figure S7.** The response of mouse fecal proteome to the presence of microbial consortiums containing fungi and/or bacteria. Statistical comparison of quantified mouse proteins (t-test, FDR 5%) was performed in the Perseus proteomic software on A) B and BY, B) B and GF, C) BY and Y, and D) GF and Y groups. Functional classes were based on the mouse protein annotations derived from the UniProtKB database[29] and compared to those obtained by the DAVID[30] and STRING-db[31] tools.

**Table S6** Number of proteins with significantly increased levels for selected bacterial strains and between different mice groups

| Bacterial Strain | Proteins Quantified | ANOVA FDR 5% | THSD | B vs BY | | BY_ABX vs BY_AFX | | BY vs BY_AFX | | BY vs BY_ABX | | B vs BY_AFX | | B vs BY_ABX | |
|---|---|---|---|---|---|---|---|---|---|---|---|---|---|---|---|
| | | | | B | BY | ABX | AFX | BY | AFX | BY | ABX | B | AFX | B | ABX |
| *Akkermansia muciniphila YL44* | 570 | 308 | 280 | 5 | **67** | 18 | 17 | 4 | **105** | 4 | **114** | 3 | **204** | 1 | **206** |
| *Muribaculum Intestinale YL27* | 393 | 233 | 206 | 6 | **20** | **48** | 15 | 7 | **98** | 0 | **122** | 22 | **109** | 4 | **179** |
| *Blautia coccoides YL58* | 502 | 271 | 244 | **51** | 1 | **44** | 13 | 5 | **142** | 4 | **157** | 7 | **156** | 10 | **127** |
| *Clostridium clostridioforme YL32* | 921 | 553 | 456 | **100** | 7 | **125** | 27 | **104** | 83 | 36 | **147** | **211** | 80 | **120** | 96 |
